# Maintaining tandem movement cohesion through antennal movements in termites

**DOI:** 10.1101/2025.02.13.638054

**Authors:** Nobuaki Mizumoto, Sam Reiter

## Abstract

How do animals coordinate their motion during migration? Traditional models of collective motion, for example, describing bird flocks or fish schools, rely on visual interactions. However, many animals are blind, requiring movement coordination through maintenance of physical contact. The risks and cost of becoming accidentally separated may encourage the evolution of compensatory strategies. Here we study tandem running in blind termites. We quantitatively investigate how these animals use their appendages to maintain stable pair movements. During tandem runs, male followers use shorter palps and longer antennae to maintain physical contact with female leaders. Our posture-tracking analysis revealed that termites dynamically change their antennal movements. Males stabilize their antennae to maintain contact with their partners while their palps are in contact. When the male palps lost contact with a female, males started swinging antennae while increasing movement speed. Antennae removal experiments revealed that antennal swinging contributes to pair maintenance, and males compensate for single antenna loss by increasing the swinging of the remaining one. By providing detailed information on contact-based movement coordination, our results contribute to understanding the diversity of animal collective behavior.

## Introduction

When groups of animals travel together, they need to coordinate to maintain cohesion (1, 2). Social interactions mediating this coordination are often pairwise and usually happen between closest neighbors, even if the group size is large (3–5). Individuals in a group modify their speed and heading directions according to the distance and relative orientations of their partners (6, 7). Models of collective behavior show that local interactions between neighbors enable groups to coordinate their motion during migrations (8). However, such systems explicitly or implicitly assume visual communication, where individuals can know inter-individual distances immediately, such as in the collective movements of fish schools, bird flocks, insect swarms, and human groups (9). Thus, little is known about movement coordination based on other communication channels. In this study, we quantitatively describe movement coordination based on physical communication, often observed in blind arthropods.

One of the modes of collective motion of terrestrial animals is alignment in a line, often described as a processionary (e.g., caterpillars and weevil) (10–14), queuing (e.g., lobster, trilobite) (15–17), and tandem running (e.g., ants and termites) (18, 19). These systems do not use visual information for movement coordination. Many do not use long-range chemical signals, instead relying heavily on physical contact during movement coordination. This contact is not very rigid, like swarm robotics with grippers (20) or people walking hand in hand. Instead, individuals keep actively “touching” each other to indicate their physical presence (21). Presumably, to support this behavior, species that move in a line often have long appendages (12, 15, 16). Such systems can work as long as they can maintain physical contact with others. However, relying only on physical contact is vulnerable to the accidental loss of the contacts. Thus, organisms need to develop a behavioral mechanism that mitigates the fluctuation of distances between individuals while moving together.

Mating pairs of termites show a characteristic movement coordination behavior called tandem running (22). During the mating seasons, a paired female and a male explore the environment together to search for a suitable nest site for colony foundation. In many species, females play leaders while males follow their partners (23). Tandem communication is heavily based on tactile communication (22, 24). They do not use visual cues for tandem runs (25). Females often have sex pheromones attracting males (26, 27). Still, the use of pheromones is limited during movement coordination, evidenced by the fact that male-male tandem runs are able to function as well as female-male tandem runs in some species (23, 28, 29). Pheromones are mainly used when a pair completely loses each other; the female leaders pause to wait for followers and emit sex pheromones, while male followers engage in a restricted search (30). Physical communication is achieved by two types of appendages: followers touch leaders’ abdomens with their antennae and maxillary pulp (24) (Figure 1). These two appendages are different in length, with the antenna being longer than palps. This means that these two could have different functions while maintaining movement coordination.

**Figure 1.**
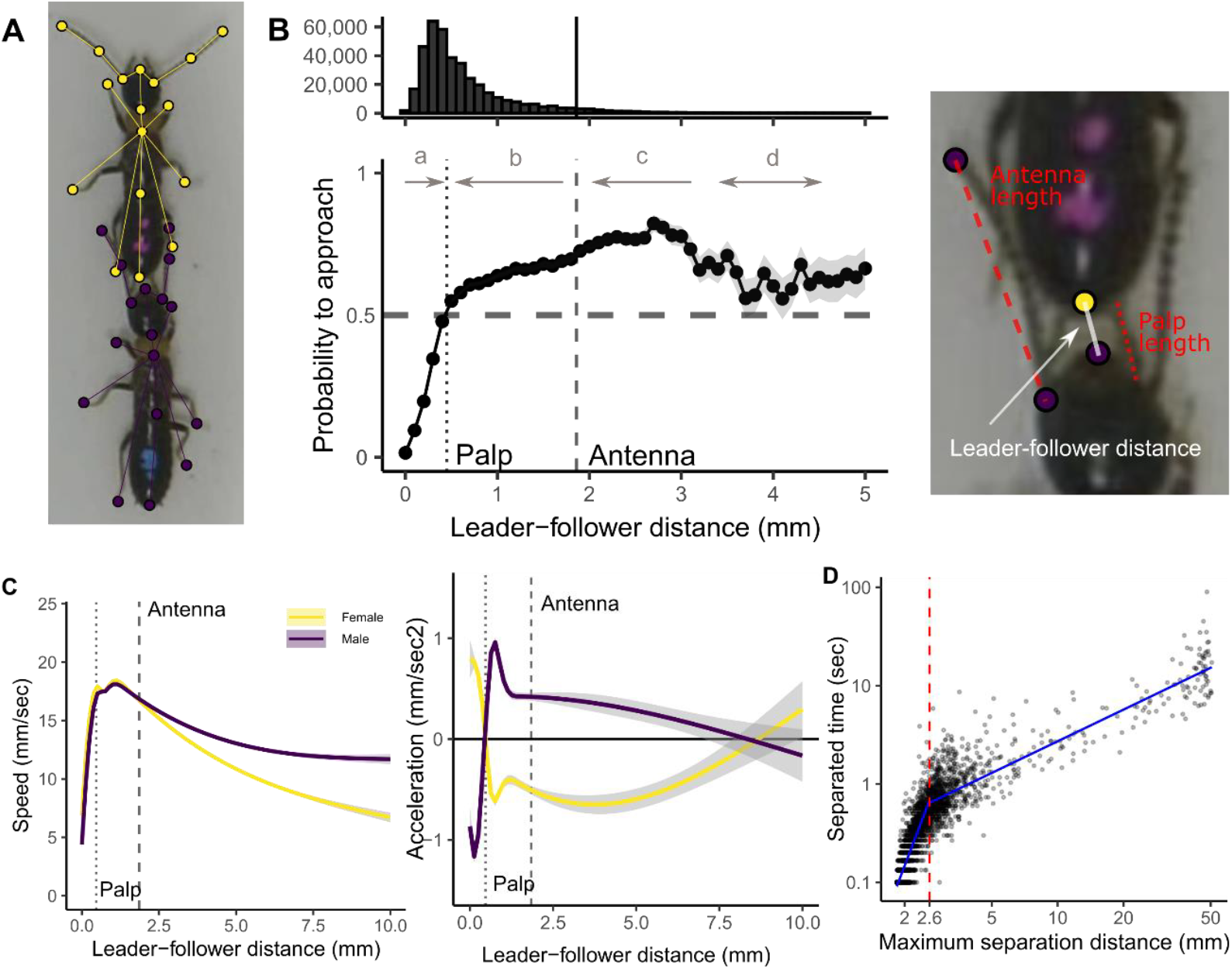
Description of tandem run by *R. speratus* in relation to appendage lengths. (A) Visualization of tracked body parts. (B) The probability that leader-follower distance becomes shorter one second later. Shaded regions indicate mean ± s.d. A dashed horizontal line indicates 0.5, above which the distance tends to get shorter one second later. The histogram shows the number of data points. (C) Termite movement speeds relative to the leader-follower distance. Lines and shaded areas indicate GAM plots with 95% confidence intervals. (D) Relationship between separation duration and maximum separated distance. Separation is defined when the leader-follower distance is larger than the antenna length. Blue lines indicate LMM regressions.

In this study, we investigate the role of antennae in termite movement coordination. First, using deep-leaning automated posture tracking (31), we quantitatively describe tandem running behavior in relation to appendage use in a termite *Reticulitermes speratus*. To further investigate the role of the antennae, we observed tandem runs of termites with the antennae removed. Finally, we compared the results with another termite species, *Coptotermes formosanus*, whose females have contact-based pheromones, compared with volatile pheromones in *R. speratus*. Our results suggest that palps work like suspension between partners, while antennae play a buffer role to prevent separation and assist stable tandem coordination.

## Methods

### Tandem observations

Alates of *Reticulitermes speratus* were collected with a piece of nesting wood from 5 colonies in Fukui and Ishikawa prefectures, Japan, in 2022 (colony name: 455, 456, 458, 459, 460), just before the swarming season. All nesting wood pieces were maintained at 22 °C until experiments. Before each experiment, we transferred nests to a room at 27°C, which promoted alates to emerge and fly. Alates were then collected, separated by sex, and color-marked with one dot of paint (PX-20; Mitsubishi) on the abdomen to distinguish sex identities. These termites were isolated individually more than 30 min before the experiments. We used individuals who shed their wings by themselves within 12 hours.

We introduced either a female-male pair, single female, or single male into the experimental arena. The experimental arenas consist of a petri dish (φ = 55 mm) covered with a layer of moistened plaster that was polished before each trial. We recorded termite movements in the arena for 15 minutes using a video camera (HC-X1500-K, Panasonic) with a resolution of 3840x2160 pix at 59.96 frames per second (FPS). In total, we obtained the videos of 20 females, 20 males (455x5, 458x5, 459x5, 460x5), and 21 female-male pairs (455x5, 458x5, 459x5, 460x5 + 456x1). Note that we only used nest-mate pairing in our experiments, which does not affect tandem running behavior in this species (23). All the videos were cropped to 2000x2000 pix to only include the arena in the frame before the video analysis.

### Posture tracking

All videos were analyzed using SLEAP v 1.4.0 (31) to estimate the movement of body parts of each individual. We used a 17-node skeleton: antenna tips (LR), antenna middle (LR), antenna base (LR), head (middle of mouth parts), head-pronotum boundary, pronotum-mesonotum boundary, metanotum-abdomen boundary, abdomen-tip, fore legs (LR), mid legs (LR), and the hind legs (LR). We labeled 354 individuals from 45 videos for training. We trained a U-Net-based model with a multi-animal top-down approach, with a receptive field size of 76 pixels for the centroid and 156 pixels for the centered instance, on Nvidia GeForce RTX 4090, where augmentation was done by rotating images from -180 to 180 degrees. The mean Average Precision (mAP) and mean Average Recall (mAR) of this model were 0.687 and 0.766, respectively. While tracking after the inference, we used the instance similarity method with the greedy matching method. All pose estimation data were converted to HDF5 files for further analysis.

### Data analysis

We used Python to format all HDF5 files for further analysis and converted them into FEATHER files for analysis in R (32). We employed a linear interpolation method to address missing values in the dataset after down-sampling data into 30 FPS. After scaling all data from pixels to mm (2000 pixels = arena size), we used a median filter with a kernel size of 5 to reduce noise.

After data formatting, we obtained measurements relating to termite movements. First, we obtained the angles of the antennae as angles between two vectors: antenna base-antenna tip and pronotum-head (Fig. 1). We estimated the antennae movement as the change of this angle between successive frames. The autocorrelation obtained for this antennae movement decayed rapidly and became negative with a lag of 1 (Fig. S1). Thus, we analyzed antenna movement by sampling it at 30 FPS. Second, as tandem running is maintained by male followers contacting female leaders with their mouthparts in the studied species, we measured the distance between the head of males and the abdomen of females (Fig. 1A). We used this following distance as an indicator of tandem running. We especially examined how the current following distance can predict the future following distance. The autocorrelation function of the following distance decayed gradually and approached 0 around lag 30 (Fig. S1). So, we used the threshold of 30 (= 1 second) as a reference point and investigated if the following distance became smaller or larger 1 second later, given the current following distance. Third, we investigated the moving speed and acceleration of termite bodies by focusing on the trajectories of pronotum. The speed was obtained by calculating the distance displacements of the pronotum, while the acceleration was obtained by calculating the time difference in speed. These measurements were obtained at 30 FPS.

We also estimated the antennae length from the posture datasets by summing up the distances from the antennae base to the antennae middle and the distance from the antennae middle to the antennae tip. The antennae length was measured for every frame and averaged for each individual. Then, we obtained the average across individuals to obtain a species-specific value of 1.86 mm for *R. speratus*. We then manually measured the length of maxillary palps for follower males for each video; the average was 0.47 mm for *R. speratus*.

We regarded pairs as within physical contact when the following distance was within the antennae length. Conversely, we analyzed each separation event as a series of frames during which the following distance was larger than the antenna length. For each separation event, we recorded the separation duration and the maximum distance between partners. The relationship between separated distance and duration was apparently different between short distance (quick catching up events) and long distance (random searching) (Fig. 1). We investigated this breakpoint as follows. First, we sequentially changed the breakpoint from 2 mm to 5 mm in 0.01 mm. Then, we assigned data a category, below or above the breakpoint. We used a linear mixed model (LMM) for the datasets, where separated duration was the response variable; separated distance, breakpoint category, and interaction were explanatory variables; each pair was included as a random effect (random intercept). We obtained AIC for each breakpoint and identified the breakpoint that minimized the AIC as the best breakpoint. Note that the AIC of the model without breakpoint was larger than this.

### Statistical analysis

We compared the degrees of antenna movement between sexes and across different contexts, including before pairing (females and males were observed isolated), during tandem runs with the following distance being within antenna length, during tandem runs with the following distance being within palp distance, and when separated after tandem running. For each individual and each context, we obtained the average angle at which termites moved their antenna per frame in radians. The right and left antennae values were averaged to get one value for each individual and context. Then, we compared this averaged antenna moving angle between sexes for each context, using LMM with the original colony included as a random effect (random intercept). We also compared antenna movement angles among these contexts within each sex, using similar LMMs, but each individual ID nested within each colony was included as a random effect.

All LMM analyses were conducted using the lmer() function in the package ‘lme4’ (33). We used the likelihood ratio (type II) test to investigate each explanatory variable’s statistical significance, using the Anova() function in the package ‘car.’ In cases of significant effects with multiple levels, we ran Tukey’s post hoc test using the glht() function in the ‘multcomp’ package (34). We also used the r.squaredGLMM() function from the ‘MuMIn’ package (35) to calculate marginal and conditional *R*^2^ values for our LMMs to obtain effect sizes.

### Antenna removal

In order to investigate further how termite followers use antennae while tandem running behavior, we cut the antennae of males and tested how a lack of antennae affects movement coordination. We used allates from 3 colonies (C, F, G) collected in Hyogo prefecture, Japan in 2023. After preparing and color-marking individuals in the above process, we placed a dish with termites on an ice pack to immobilize them. Under a stereomicroscope, the right antenna was clipped off either at the base of the head (full cut) or the middle (half cut). As controls, we prepared termites that were immobilized but did not receive antennae cut (control) and that received both antennae cut at the base of the head (no antennae). We isolated each termite for 30 minutes after antenna removal. Then, we introduced one treated male and an untreated female to the observational arena to record their behavior as above. In total, we obtained videos of 17 no antennae (Cx6, Fx5, Gx6), 15 full-cut (Cx5, Fx4, Gx6), 13 half-cut (Fx6, Gx7), and 18 control (Cx7, Fx5, Gx6). These videos were analyzed using the same approach as above. Using SLEAP, we labeled 58 individuals from 12 videos for training. We trained the model based on the model developed above by resuming the training, where the mAP and mAR were 0.57 and 0.7, respectively.

We used data from pairs that showed tandem runs for more than 50% of the observational period for antenna removal experiments. We compared antenna movements among treatments (control, half-cut, full-cut) when termites are in tandem (within antenna distance) or after separation. Because the antenna removal treatment was always the right antenna, we compared the movement of the left antenna. We used LMMs with the original colony included as a random effect (random intercept). We also examined the relationship between antenna movement and time spent in tandem runs across all pairs using a generalized linear mixed model (GLMM), where antennae movement, antennae treatment, and their interactions were included as explanatory variables, while the original colony was included as a random effect (random intercept).

### Interspecific comparison

We performed the same tandem description in *Coptotermes formosanus* as a comparison. Alates of *C. formosanus* were collected using light traps during a dispersal flight in Okinawa, Japan, on May 22nd, 2022. Because *C. formosanus* shows synchronized flight among colonies within the same area, it is presumed that alates originated from multiple colonies. We used individuals who shed their wings by themselves within 12 hours. We used the experimental arenas made of larger dishes (φ = 90 mm) due to their larger body size than *R. speartus*. We obtained the videos of 10 females, 10 males, and 10 pairs. We estimated palp length and antenna length as 0.62 mm and 2.25 mm, respectively.

We analyzed the data of *C. formosanus* in the same way as *R. speratus* described above. We also compared the duration of tandem running (the time while following distance of partners was within the antenna length) using the mixed-effects Cox model (coxme() function in the ‘coxme’ package in R (36)), with species as a fixed effect and pair id as a random effect.

## Results

### Anatomy of tandem running in relation to appendages

Our posture tracking revealed that termites adjust behaviors and inter-individual distances based on the appendage contacts between leaders and followers. Paris appeared to maintain palp length distance between themselves (median inter-individual distance = 0.47mm, mean palp length = 0.47mm), where 50% of data comes between 0.32 to 0.77mm (Fig. 1B). When the distance is closer than palp length, the pair tended to move slightly apart (a in Fig. 1B). On the other hand, pairs approached each other when they lost contact with palps (b in Fig. 1B). These dynamics were achieved through speed control. When the palp contacted, the movement speed of the pair was slower; the leader sped up, and the follower slowed down (Fig. 1C). On the other hand, tandem running pairs moved at the maximum speed when the palps lost contact, but while the partner was within the range of the antenna. Once palp contact was lost, males sped up to catch up with females, and females slowed down. This speed regulation was stronger in males than females, indicating males more actively adjust their movement to coordinate motion with females. These observations are consistent with previous studies on descriptive observations (24) or analyses focusing on the body centers for tracking (23, 25, 37). However, our analyses quantitatively show that termite movement coordination is clearly regulated by tactile communications based on the touching ranges of their appendages.

Interestingly, pairs recovered contact after a separation event (c in Fig. 1B). Among 2,491 separation events, pairs came back to the range of antennal distance within 1 sec in 92.5% of events. Similarly, pairs separated not more than two antennal distance (3.69 mm) in 91.6% of events. This is due to their movement coordination, where male speeds exceeded female speeds once they lost antennal contacts (Fig 1C). This is known as the efficient reunion strategy (30, 38). However, once they were further separated, they got lost, and interactions became stochastic (d in Fig. 1B). Our analysis of the maximum separated distance and time required for reunion revealed that there were two different types of separation events. When the separated distance was short, the required time was sensitive to the separated distance, reflecting a short-term catching-up process. On the other hand, once pairs were separated by further distance, this relationship was relaxed, reflecting a long-term searching process. The threshold was relatively small (2.6mm), indicating that termites do not use distance communication methods (e.g., vision or olfactory) during the tandem running process.

### Antenna use during tandem runs

Antenna movement patterns were clearly associated with different phrases of tandem running in both females and males, where we found sex differences in each phase (LMM, *P* < 0.05) and across phases (Fig. 2AD). Before pairing with a mating partner, both females and males actively moved to look for a mating partner (Fig. 2A) (30). During this phase, both females and males moved their antennae while sensing the environments, where males moved their antennae more than females, albeit with a small effect size (LMM, χ^2^_1_ = 4.30, *P* = 0.038, *R*^2^_m_ = 0.09, Fig. 2CD). When the pair was tandem running, leader females moved their antennae to sense the environments, searching for potential nest sites (Fig. 2BC). On the other hand, male followers dynamically changed their antennae movements. Males often stabilized their antennae with lower angles by touching the leaders’ body side (Fig. 1AB, Fig. 2C). This was prominent when the tandem pair was within palp length distance, meaning that followers could physically contact leaders with two types of appendages (palps and antennae) (Fig. 2D). Because of this antennae stabilization, male antenna movement was smaller than females (LMM, within-antenna: χ^2^_1_ = 10.4, *P* = 0.001, *R*^2^_m_ = 0.23, within palp: χ^2^_1_= 66.7, *P* < 0.001, *R*^2^_m_= 0.65, Fig. 2CD). Once they lost contact with their palps, male followers began to move antennae, where antennae movements were larger (stat, Fig. 2BD). When termites lost physical contact at all (the distance between pairs was larger than antennae length), males moved their antennae as in searching for a partner before pairing (Fig. 2BCD), while waiting females slowed down their antennae movements (LMM, χ^2^_1_= 19.7, *P* < 0.001, *R*^2^_m_ = 0.39, Fig. 2BD).

**Figure 2.**
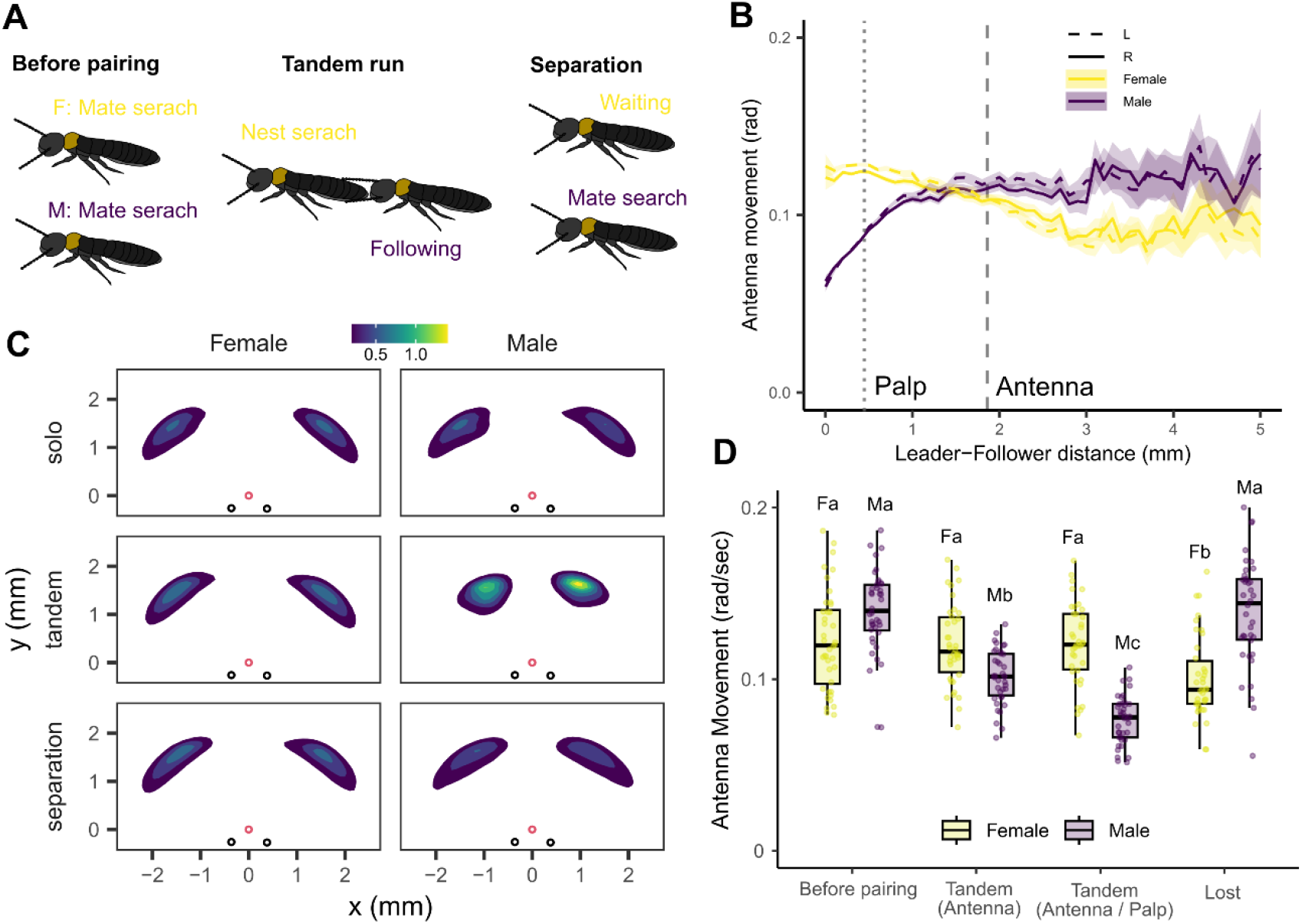
Antennae movement during tandem pairing in *R. speratus*. (A) Behavioral phases of tandem pairing. Before females and males encounter a partner, both sexes actively search for a mating partner. Once they encounter, female leaders look for a nest site, while males follow females. When they are separated, males engage in local intensive search, while females pause to wait for partners. (B) Antennae movement in relation to female-male distances. Shaded regions indicate mean ± s.d. (C) The relative position of antennae tips. Red circles indicate the position of the head tip, while black circles indicate the positions of antenna bases. (D) Comparison of mean antennae movements between sexes and across tandem pairing status. Different letters indicate significant differences within sex comparison (LMM, *P* < 0.05).

### Compensation for antenna loss

Termites lacking antennae had limited following ability. First, males without antennae at all did not show any following behavior (0/17), where a few pairs show occasional reversed tandem running with the female following the male (2/17). The reversed tandem run should be technically possible as females show female-female tandem runs in this species (23, 29), although it has never been observed in many efforts of tandem observations on this species (23, 25, 29, 30, 39). Note that these reversed tandem runs were less functional because males without antennae were less active than other treatment males (male moved distance, LMM, χ^2^_3_ = 17.2, *P* < 0.001; Tukey HSD, *P* < 0.05 for all combinations between without antenna and others). As a result, reversed tandem running did not move for long distances; even the longest reversed tandem running event ended after the leader moved 80 mm, while the longest runs were 251 mm (one antenna), 444 mm (half-cut antennae), and 1,062 mm (control), respectively. However, we did not observe a significant difference between treatments in the distance of the longest tandem running events for each pair (LMM, χ^2^_3_ = 3.32, *P* = 0.35) with the limited sample size.

In males without one antenna, only two pairs showed tandem runs for more than 50% of the observation period (2/15; Fig. 3A). Males with a half-cut antenna showed tandem runs for more than 50% of the observation period in 3/13 pairs, while control males did in 8/18 pairs (Fig. 3A). Note that this level was lower than the observation of tandem runs above (16/20), suggesting that icing treatment reduced performance of tandem runs. Among these pairs, there was positive relationship between the proportion of time spent in tandem runs and antennae movements when termites got separated (GLMM, *P* < 0.001, Fig. 3A). The relationship between leader-follower distance and antenna movements was consistent with that of natural tandem runs, among pairs that showed tandem runs for > 50% of time (Fig. 3B). However, termites with trimmed antennae moved their antennae more than intact individuals. When followers lost contact with females with their antennae, males with only a single antenna moved their antenna more than that of intact males (LMM, χ^2^_2_ = 13.4, *P* = 0.001, *R*^2^_m_ = 0.29, Fig. 3A). This increased antenna activity should have contributed to tandem maintenance.

**Figure 3.**
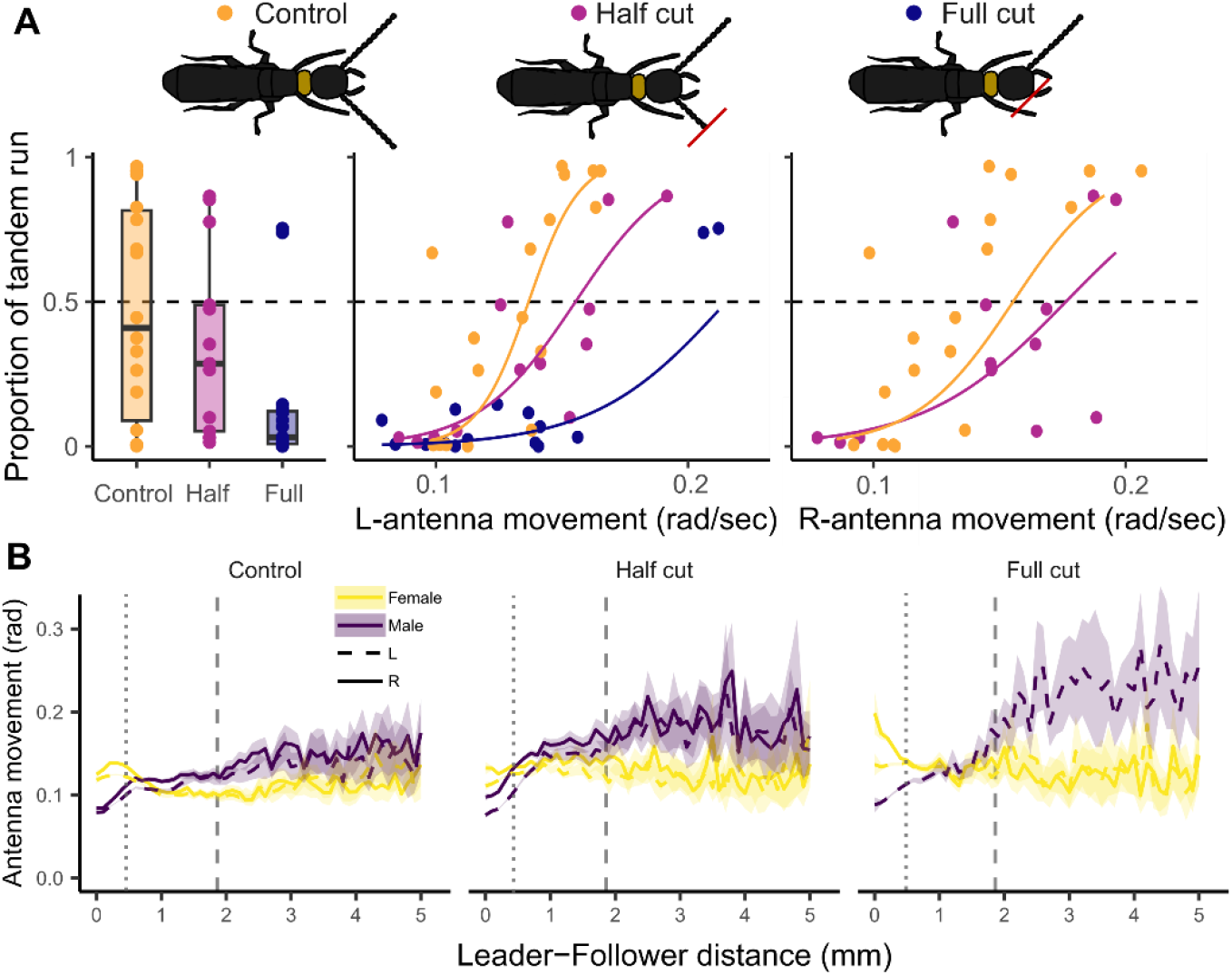
The effect of antenna removal on tandem coordination. (A) Time spent in tandem running for males with antenna treatments. The solid lines are the regression curve calculated with GLMM, indicating significant positive relationships. (B) Antenna movements in relation to female-male distances. Shaded regions indicate mean ± s.d. Vertical lines indicate the length of palps and antennae.

### Interspecies comparison

Our observation on *C. formosanus* showed overall similar patterns to *R. speratus*, where the pair also changed movement coordination and antenna motion according to the inter-individual distance relating to appendage lengths (Fig. 4). Different from *R. speratus*, the pairs kept a shorter distance between leaders and followers than palp length (50% of data between 0.24 to 0.63mm, and mean palp length = 0.62, Fig. 4A). This was because the male head and the female abdomen tip often vertically overlapped in *C. formosanus* (Fig. 4A), while females and males were usually at the same height in *R. speratus* (Fig. 1A). Movement speed was highest around palp length (Fig. 4B); once pairs lost contact with palps, males accelerated, and females accelerated to maintain contact (Fig. 4BC). The tandem duration (pair within the antenna length) was not different between *C. formosanus* and *R. speratus* (mixed-effects Cox model, χ^2^_1_ = 0.21, *P* = 0.645). When they lost physical contact, a pair returned to the antennal distance range within 1 sec in 86.6% of events and within the two antennal distances (4.51 mm) in 73.2% of events. Similar to *R. speratus*, there were two types of separation events: short-term catching-up and long-term searching (Fig. 4D).

**Figure 4.**
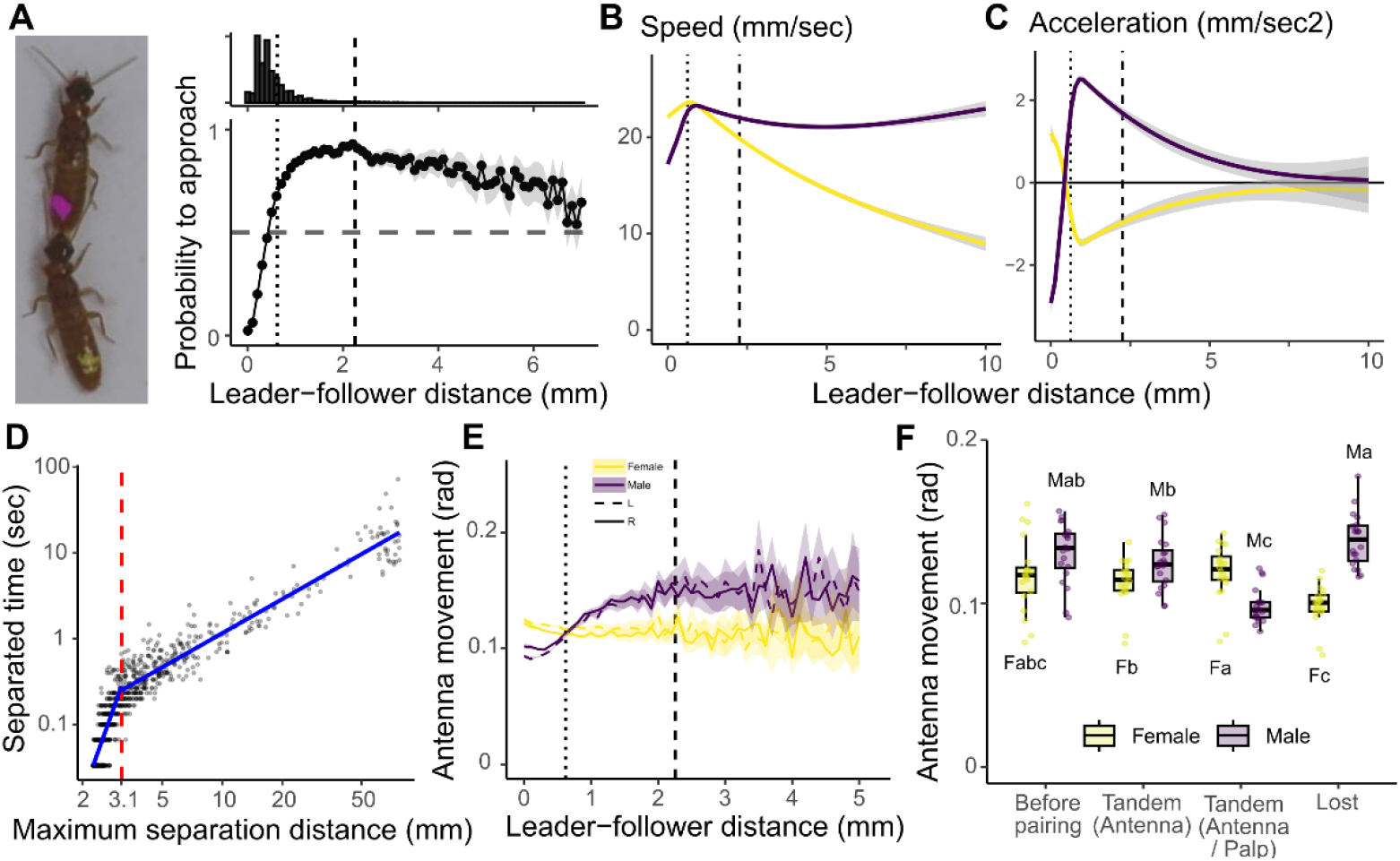
Analysis of *C. formosanus*. (A) The probability that leader-follower distance becomes shorter one second later. Shaded regions indicate mean ± s.d. A dashed horizontal line indicates 0.5, above, which means the distance tends to get shorter one second later. The histogram shows the number of data points. Dashed vertical lines indicate palp and antenna length, respectively. (B-C) Termite movement speeds relative to the leader-follower distance. Lines and shaded areas indicate GAM plots with 95% confidence intervals. (D) Relationship between separation duration and maximum separated distance. Separation is defined as when the leader-follower distance is larger than the antenna length. Blue lines indicate LMM regressions. (E) Antennae movement in relation to female-male distances. Shaded regions indicate mean ± s.d. (F) Comparison of mean antennae movements between sexes and across tandem pairing status. Different letters indicate significant differences within sex comparison (LMM, *P* < 0.05).

*C. formosanus* also used antennae differently according to the phase of tandem running, where searching individuals moved antennae actively while following males and waiting females reduced antenna movements (Fig. 4EF). Before pairing, both females and males moved their antennae while sensing the environments without sex differences (LMM, χ^2^_1_ = 2.28, *P* = 0.131, *R*^*2*^_*m*_ = 0.10, Fig. 4F). During tandem runs, females moved their antennae to search for nest sites, while male followers stabilized their antennae by touching leaders’ body side while maintaining palp contact (LMM, χ^2^_1_ = 34.2, *P* < 0.001, *R*^*2*^_*m*_ = 0.34, Fig. 4AEF). Once they lost palp contact, male followers began to move their antennae, where antennae movements were larger than females but with a small effect size (LMM, χ^2^_1_ = 10.6, *P* = 0.001, *R*^*2*^_*m*_ = 0.13, Fig. 4EF). When termites totally lost physical contact, males moved their antennae as in searching for a partner before pairing, while waiting females slowed down their antennae movements (LMM, χ^2^_1_ = 195.1, *P* < 0.001, *R*^*2*^_*m*_ = 0.66, Fig. 4EF).

## Discussion

In blind animals unable to communicate at a distance, partners need to keep physical contact to maintain cohesion during movement coordination. We showed that tandem running termites use two appendages (palp and antenna) to loosely maintain physical contact during movement coordination, with partners adjusting their movements according to sensing status (Fig. 1, 4). First, termites use palps to maintain a desired distance between partners, accelerating or decelerating their movements according to palp contact. In this sense, palps work like a suspension system coupling two trains, where they are important for smooth operation but not essential for moving together. This function is consistent with previous observations that removing palps has a smaller impact than removing antennae for tandem running (24). Second, antennae play an essential role in movement coordination for active sensing, as males without antennae cannot follow females. In contact ranges, followers stabilize their antennae to ensure continued contact. Followers then actively move antennae as they are separated for searching and sensing. These results indicate that longer appendages lead to better movement coordination, explaining that many species with contact-based collective motion have long appendages (12, 15, 16).

Among blind insects, thigmotaxis is a commonly used movement mechanism based on tactile information (40–42). Cockroaches are known to use antennae for active tactile sensing to localize and move toward objects (43, 44). During this process, cockroaches keep touching the object while moving towards it. When an object is in front of the animal, they keep it between two antennae during approach (43, 44). Also, when cockroaches exhibit wall-following behavior, they use their antennae to sense and maintain their distance from the wall (45–47). Our study suggests that termite followers use a mechanism similar to these observations on thigmotaxis for tandem running. Male followers used antennae during tandem runs by touching female followers to maintain distance. This mechanism can tell the movement direction of the leader; if a sharp turn of the leader results in the loss of one antenna contact, the follower can recognize the turn direction by proceeding in the direction where antenna contact is maintained. Given that thigmotaxis and wall-following are widely observed across a wide range of cockroaches (48), including most termite lineages (49–51), termites might evolutionary reuse the wall-following behavioral mechanism as a preadaptation trait for tandem running coordination.

Our comparison between *R. speratus* and *C. formosanus* shows that these species use tactile communication similarly (Fig. 1, 4). This suggests they rely on contact signals and do not use long-distance communication during movement coordination. In *Reticulitermes* termites, when a pair is completely separated, female leaders pause for a long period (30) and engage in calling behavior, perhaps to emit sex pheromones to attract stray male partners (52, 53). In contrast, the sex pheromones of *C. formosanus* work as contact pheromones (24, 54), and females do not pause for long compared to *R. speratus* upon accidental separation (30) with limited calling behavior (24, 54). Accordingly, a previous study within a larger arena showed that it took longer for *C. formosanus* than *R. speratus* to reunite once they were accidentally separated (30). Note that we did not observe this difference in this study, perhaps due to the smaller arena size necessary to obtain higher-quality videos. Thus, these two species use distinct behaviors for communicating with a partner at a longer distance, contrasting with similar coordinated mechanisms at a shorter distance.

Our results also highlight similarities and differences between visual-based movement coordination and tactile-based tandem coordination. For example, behavioral models of vision-based collective motion often consider long-range attraction and short-range repulsion described by the interactive zones with different radiuses (55–57). In these models, agents interact with neighbors quantitatively based on the estimation of pairwise distances between neighboring individuals, using long-range signaling mechanisms. In the case of termites, although they do not use vision for communications, we found that they have multiple regions similar to long-range attraction and short-range repulsion; when partners are within the palp distance, they increase distance, while when partners are more than palp but within antenna distance, they shorten their distance. Termite distance estimation mechanisms are likely distinct from those of visual communication, as shown in a physical model for robotic antennae (45, 58), requiring further work on models of contact-based movement coordination.

Tandem running behavior looks like easy teamwork; two individuals simply move together by maintaining physical contact. However, closer inspection reveals that it is built from fine-scale movement regulation, where leaders and followers change their movement patterns according to their physical contact state. A theoretical understanding of such movement coordination is lacking compared to other forms of collective behavior. Although several studies modeled tandem running recruitment in ants using simulation (59–62) or the kilobot platform (63), these studies focused on colony-scale recruitment dynamics rather than tandem running behavior or movement coordination. Robot-insect interactions may offer a promising direction for studying this process in greater detail (64), likely requiring further fine-tuning of robot behavior according to insect models. Our study provides baseline information on contact-based movement coordination through quantifying termite tandem runs, which could be expanded to multiple systems through modeling.

## Data accessibility

The data and codes are available on GitHub: nobuaki-mzmt/Tandem_antenna_role.

## Acknowledgment

We thank Aoi Mizumoto for the help during the video analysis, Kensei Kikuchi for help during sampling termites, and Dr. Thomas Bourguignon for providing experimental spaces. We also thank Sayomi Kamimoto and the members of the computational neurology unit at OIST for helpful discussion. This study is supported by a JSPS (Japan Society for the Promotion of Science) Research Fellowship for Young Scientists CPD (Cross-border Post Doctorate) (20J00660) to N.M., a Grant-in-Aid for Early-Career Scientists (21K15168) to N.M., IPSF fellowship from OIST to N.M., OIST core funding, and USDA National Institute of Food and Agriculture, Hatch projects number 7007938.

## Author contributions

N.M.: conceptualization, data curation, formal analysis, funding acquisition, investigation, methodology, project administration, validation, visualization, writing-original draft S.R.: conceptualization, methodology, supervision, writing-review & editing

## Notes

### Competing Interest Statement

The authors have declared no competing interest.

## References

1. I. D. Couzin, J. Krause, Self-organization and collective behavior in vertebrates. Advances in the Study of Behavior 32, 1–75 (2003).

2. D. Gorbonos, et al., An effective hydrodynamic description of marching locusts. Phys. Biol. 21, 026004 (2024).

3. M. Ballerini, et al., Interaction ruling animal collective behavior depends on topological rather than metric distance: evidence from a field study. Proceedings of the National Academy of Sciences of the United States of America 105, 1232–1237 (2008).

4. I. Ashraf, et al., Simple phalanx pattern leads to energy saving in cohesive fish schooling. Proceedings of the National Academy of Sciences 114, 9599–9604 (2017).

5. L. Lei, R. Escobedo, C. Sire, G. Theraulaz, Computational and robotic modeling reveal parsimonious combinations of interactions between individuals in schooling fish. PLoS Computational Biology 16, e1007194 (2020).

6. Y. Katz, K. Tunstrøm, C. C. Ioannou, C. Huepe, I. D. Couzin, Inferring the structure and dynamics of interactions in schooling fish. Proceedings of the National Academy of Sciences of the United States of America 108, 18720–18725 (2011).

7. J. E. Herbert-Read, Understanding how animal groups achieve coordinated movement. Journal of Experimental Biology 2971–2983 (2016). 10.1242/jeb.129411.

8. S. Camazine, et al., Self-organization in Biological Systems (NJ: Princeton University Press, 2001).

9. N. T. Ouellette, A physics perspective on collective animal behavior. Phys. Biol. 19, 021004 (2022).

10. M. J. Steinbauer, Thigmotaxis maintains processions of late-instar caterpillars of Ochrogaster lunifer. Physiological Entomology 34, 345–349 (2009).

11. P. Collard, Processionary Caterpillars at the Edge of Complexity. Artificial Life 30, 171–192 (2024).

12. T. D. Fitzgerald, A. Pescador-Rubio, The Role of Tactile and Chemical Stimuli in the Formation and Maintenance of the Processions of the Social Caterpillar Hylesia lineata (Lepidoptera: Saturniidae). Journal of Insect Behavior 15, 659–674 (2002).

13. M. Uemura, L. E. Perkins, M. P. Zalucki, A. Battisti, Movement behaviour of two social urticating caterpillars in opposite hemispheres. Mov Ecol 8, 4 (2020).

14. T. D. Fitzgerald, A. Pescador-Rubio, M. T. Turna, J. T. Costa, Trail Marking and Processionary Behavior of the Larvae of the Weevil Phelypera distigma (Coleoptera: Curculionidae). Journal of Insect Behavior 17, 627–646 (2004).

15. W. Herrnkind, Queuing Behavior of Spiny Lobsters. Science 164, 1425–1427 (1969).

16. J. Vannier, et al., Collective behaviour in 480-million-year-old trilobite arthropods from Morocco. Scientific Reports 9, 1–10 (2019).

17. X. G. Hou, D. J. Siveter, R. J. Aldridge, D. J. Siveter, Collective behavior in an early Cambrian arthropod. Science 322, 224 (2008).

18. E. L. Franklin, The journey of tandem running: The twists, turns and what we have learned. Insectes Sociaux 61, 1–8 (2014).

19. N. Mizumoto, C. R. Reid, Ant and termite collective behavior: Group-level similarity arising from individual-level diversity. Ecological Research 1440-1703.12510 (2024). 10.1111/1440-1703.12510.

20. V. Trianni, M. Dorigo, Self-organisation and communication in groups of simulated and physical robots. Biol Cybern 95, 213–231 (2006).

21. T. J. Prescott, M. E. Diamond, A. M. Wing, Active touch sensing. Philosophical Transactions of the Royal Society B: Biological Sciences (2011). 10.1098/rstb.2011.0167.

22. W. L. Nutting, “Flight and colony foundation.” in Biology of Termites, K. Krishna, F. M. Weesner, Eds. (Academic Press, 1969), pp. 233–282.

23. N. Mizumoto, T. Bourguignon, N. W. Bailey, Ancestral sex-role plasticity facilitates the evolution of same-sex sexual behavior. Proceedings of the National Academy of Sciences of the United States of America 119, e2212401119 (2022).

24. A. K. Raina, J. M. Bland, J. C. Dickens, Y. I. Park, B. Hollister, Premating behavior of dealates of the Formosan subterranean termite and evidence for the presence of a contact sex pheromone. Journal of Insect Behavior 16, 233–245 (2003).

25. N. Mizumoto, T. Bourguignon, Light alters activity but does not disturb tandem coordination of termite mating pairs. Ecological Entomology een.13209 (2022). 10.1111/een.13209.

26. C. Bordereau, J. M. Pasteels, “Pheromones and chemical ecology of dispersal and foraging in termites” in Biology of Termites: A Modern Synthesis, D. E. Bignell, Y. Roisin, N. Lo, Eds. (Springer Netherlands, 2011), pp. 279–320.

27. Y. Mitaka, T. Akino, A review of termite pheromones: multifaceted, context-dependent, and rational chemical communications. Frontiers in Ecology and Evolution 8, 595614 (2021).

28. G. Li, X. Zou, C. Lei, Q. Huang, Antipredator behavior produced by heterosexual and homosexual tandem running in the termite Reticulitermes chinensis (Isoptera: Rhinotermitidae). Sociobiology 60, 198–203 (2013).

29. K. Matsuura, E. Kuno, T. Nishida, Homosexual Tandem Running as Selfish Herd in Reticulitermes speratus: Novel Antipredatory Behavior in Termites. Journal of Theoretical Biology 214, 63–70 (2002).

30. N. Mizumoto, S. Dobata, Adaptive switch to sexually dimorphic movements by partner-seeking termites. Science Advances 5, eaau6108 (2019).

31. T. D. Pereira, et al., SLEAP: A deep learning system for multi-animal pose tracking. Nature Methods 19, 486–495 (2022).

32. R Core Team, R: A language and environment for statistical computing. (2023). Deposited 2023.

33. D. M. Bates, M. Maechler, Package “lme4” Linear Mixed-Effects Models using “Eigen” and S4. Journal of Statistical Software · (2015).

34. T. Hothorn, Multcomp: Simultaneous inference in general parametric models. R package version 1, 4 (2020).

35. K. Bartoń, MuMIn: Multi-Model Inference. (2024). Deposited 22 June 2024.

36. T. M. Therneau, coxme: mixed effects Cox models. (2015). Deposited 2015.

37. G. Valentini, N. Mizumoto, S. C. Pratt, T. P. Pavlic, S. I. Walker, Revealing the structure of information flows discriminates similar animal social behaviors. eLife 9, e55395 (2020).

38. N. Mizumoto, A. Rizo, S. C. Pratt, T. Chouvenc, Termite males enhance mating encounters by changing speed according to density. Journal of Animal Ecology 89, 2542–2552 (2020).

39. N. Mizumoto, N. Nagaya, R. Fujisawa, Wasted Efforts Impair Random Search Efficiency and Reduce Choosiness in Mate-Pairing Termites. The American Naturalist 000–000 (2024). 10.1086/732877.

40. J. Basil, D. Sandeman, Crayfish (Cherax destructor) use Tactile Cues to Detect and Learn Topographical Changes in Their Environment. Ethology 106, 247–259 (2000).

41. E. R. Hunt, et al., Ants show a leftward turning bias when exploring unknown nest sites. Biology Letters 10, 20140945 (2014).

42. J. Okada, Cockroach antennae. Scholarpedia 4, 6842 (2009).

43. J. Okada, Y. Toh, Active tactile sensing for localization of objects by the cockroach antenna. J Comp Physiol A 192, 715–726 (2006).

44. J. Okada, Y. Toh, The role of antennal hair plates in object-guided tactile orientation of the cockroach (Periplaneta americana). J Comp Physiol A 186, 849–857 (2000).

45. J.-M. Mongeau, A. Demir, J. Lee, N. J. Cowan, R. J. Full, Locomotion- and mechanics-mediated tactile sensing: antenna reconfiguration simplifies control during high-speed navigation in cockroaches. Journal of Experimental Biology 216, 4530–4541 (2013).

46. J. M. Camhi, E. N. Johnson, High-frequency steering maneuvers mediated by tactile cues: antennal wall-following in the cockroach. Journal of Experimental Biology 202, 631–643 (1999).

47. N. J. Cowan, J. Lee, R. J. Full, Task-level control of rapid wall following in the American cockroach. Journal of Experimental Biology 209, 1617–1629 (2006).

48. W. J. Bell, L. M. Roth, C. A. Nalepa, Cockroaches Ecology, Behavior and Natural History (JHU Press, 2007).

49. H. Shimoji, N. Mizumoto, K. Oguchi, S. Dobata, Caste-biased locomotor activities in isolated termites. Physiological Entomology 45, 50–59 (2019).

50. O. Miramontes, O. DeSouza, L. R. Paiva, A. Marins, S. Orozco, Lévy flights and self-similar exploratory behaviour of termite workers: beyond model fitting. PloS one 9, e111183 (2014).

51. L. R. D. Paiva, et al., Scale-free movement patterns in termites emerge from social interactions and preferential attachments. 1–39 (2020). 10.1073/pnas.2004369118/-/DCSupplemental.y.

52. F. M. Weesner, The Biology of Colony Foundation in <i>Reticulitermes hesperus<i> Banks. University of California Publications in Zoiiwgy 61, 253–314 (1956).

53. H. Buchli, Les tropismes lors de la pariade des imagos de Reticulitermes lucifugus R. (I). Vie Milieu 11, 308–315 (1960).

54. T. Chouvenc, D. Sillam-Dussès, A. Robert, Courtship behavior confusion in two subterranean termite species that evolved in allopatry (Blattodea, Rhinotermitidae, Coptotermes). Journal of Chemical Ecology 46, 461–474 (2020).

55. I. Aoki, A simulation study on the schooling mechanism in fish. Bulletin of the Japanese Society of Scientific Fisheries 48(8), 1081–1088 (1982).

56. C. W. Reynolds, Flocks, herds and schools: A distributed behavioral model. ACM SIGGRAPH Computer Graphics 21, 25–34 (1987).

57. I. D. Couzin, J. Krause, R. James, G. D. Ruxton, N. R. Franks, Collective memory and spatial sorting in animal groups. Journal of Theoretical Biology 218, 1–11 (2002).

58. J.-M. Mongeau, et al., Mechanical processing via passive dynamic properties of the cockroach antenna can facilitate control during rapid running. Journal of Experimental Biology 217, 3333–3345 (2014).

59. N. Goy, S. M. Glaser, C. Grüter, The adaptive value of tandem communication in ants: Insights from an agent-based model. Journal of Theoretical Biology 526, 110762 (2021).

60. F. Abramov, et al., Tandem Running Algorithm with Route Selection in 2024 IEEE 7th International Conference on Actual Problems of Unmanned Aerial Vehicles Development (APUAVD), (2024), pp. 298– 303.

61. F. Abramov, et al., Robot Teamwork: Active and Passive Search Algorithms. Separate Issues of Management in Swarm Robotics in 2024 IEEE 17th International Conference on Advanced Trends in Radioelectronics, Telecommunications and Computer Engineering (TCSET), (2024), pp. 454–457.

62. F. Abramov, O. Andreieva, O. Andreiev, T. Diachenko, I. Maksymenko, Adaptation of the Tandem Running Algorithm to Control a Robot Swarm in 2022 IEEE 3rd KhPI Week on Advanced Technology (KhPIWeek), (2022), pp. 1–4.

63. N. Ellis, et al., Testing the reality gap with kilobots performing two ant-inspired foraging behaviours. [Preprint] (2024). Available at: http://biorxiv.org/lookup/doi/10.1101/2024.06.02.596655 [Accessed 11 June 2024].

64. N. R. Franks, J. A. Podesta, E. C. Jarvis, A. Worley, A. B. Sendova-Franks, Robotic communication with ants. Journal of Experimental Biology 225, jeb244106 (2022).

