## Supplementary material for "Maintaining tandem movement cohesion through antennal movements in termites": Fig. S1

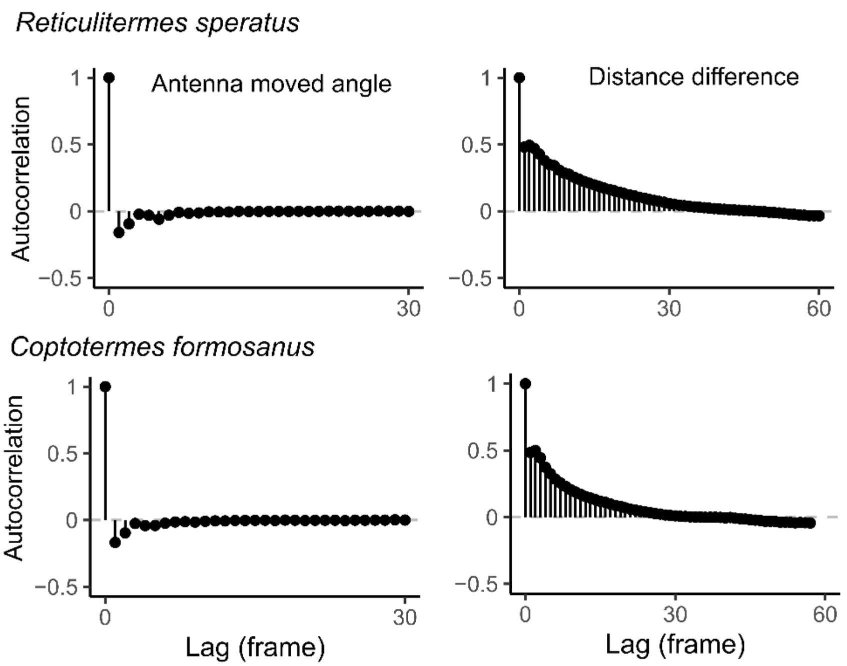


**Figure S1**. Autocorrelation function of antennae movement angles and the change in inter-individual distances.
